# Vector and cell-culture passaging of dengue clinical samples for virus isolation and amplification does not significantly change genome consensus or frequencies of intra-host viral variants

**DOI:** 10.1101/2020.08.17.254417

**Authors:** Christian K. Fung, Tao Li, Simon Pollett, Maria Theresa Alera, In-Kyu Yoon, Jun Hang, Louis Macareo, Anon Srikiatkhachorn, Damon Ellison, Alan L. Rothman, Stefan Fernandez, Richard G. Jarman, Irina Maljkovic Berry

## Abstract

Intra-host single nucleotide variants (iSNVs) have been increasingly used in genomic epidemiology to increase phylogenetic resolution and reconstruct fine-scale outbreak dynamics. These analyses are preferably done on sequence data from direct clinical samples, but in many cases due to low viral loads, there might not be enough genetic material for deep sequencing and iSNV determination. Isolation of the virus from clinical samples with low passage number increases viral load, but to date, no studies have investigated how dengue virus (DENV) culture isolation from a clinical sample impacts the consensus sequence, and there is no information on the intra-host virus population changes that may result from viral isolation. In this study, we investigate consensus and iSNV frequency differences between DENV sequenced directly from clinical samples and their corresponding low-passage isolates. Twenty five DENV1 and DENV2 positive sera and their corresponding viral isolates (*T.splendens* inoculation and C6/36 passage) were obtained from a prospective cohort study in the Philippines. These were sequenced on MiSeq with minimum nucleotide depth of coverage of 1000x, and iSNVs were detected using LoFreq. For both DENV1 and DENV2, we found that the nucleotide call concordance (including called iSNVs with variant cutoff at 5%) between direct sera sample and its cultured virus was on average 99.99%. There were a maximum of one consensus nucleotide difference between clinical sample and isolate. Interestingly, we found that iSNV frequencies were also largely preserved between the samples, with an average difference in minor variant frequency of 6.8% (95CI: 3.6%-10%) and 9.6% (95CI: 7%-12.2%) for DENV1 and DENV2, respectively. Furthermore, we found no significant differences in either DENV1 or DENV2 between the sample pairs (clinical sample and isolate) in their number of iSNV positions per genome, or in the difference in variant frequencies (*p*=0.36 and *p*=0.13, respectively, F-test). Our results show that low-passage DENV isolates may be used for identification of the majority of their human-derived within-host variant populations, which are increasingly being used for precision tracking of DENV and other RNA viruses.

## INTRODUCTION

Dengue virus (DENV) is the etiologic agent of dengue fever (DF) as well as the more severe forms of illness, dengue hemorrhagic fever (DHF) and dengue shock syndrome (DSS), which cause significant health problems each year. Approximately 50-100 million symptomatic DENV infections occur each year, with the greatest burden in tropical and subtropical regions. DENV is an arbovirus, a mosquito-borne RNA virus commonly transmitted by *Aedes* mosquitoes. It belongs to the *Flavivirus* genus of the *Flaviviridae* family and has four serotypes, DENV1, 2, 3 and 4. DENV consists of a genome approximately 11kb nucleotides (nt) in length. The genome encodes three structural proteins, nucleocapsid (C), precursor membrane (prM) and envelope (E), as well as seven nonstructural proteins (NS1, NS2A, NS2B, NS3, NS4A, NS4B, and NS5) [1–4].

Viral genome sequence data, including that of DENV, have increasingly been used in epidemics and outbreaks to provide more precise reconstructions of transmission dynamics and complement conventional non-genomic epidemiologic data. This approach of genomic epidemiology for DENV has played a crucial role in analyses of spatial and temporal outbreak dynamics, virus dispersal tracking, genotype-phenotype associations, vaccine effectiveness, vector adaptation, and many others [5–8]. Generally, genomic epidemiology studies analyze DENV consensus genomes derived from each host. However, DENV and other RNA viruses exist within a host as a population of intra-host variants, not as a single consensus genome. Advanced Next Generation Sequencing (NGS) technology can now detect intra-host viral variants present at low frequencies within a sample. Increasingly, the importance of these intra-host single nucleotide variants (iSNVs), and the amount of diversity they create within a host, has been highlighted in recent studies. For instance, Ko et al. described the presence and emergence of DENV intra-host variants with differing selection advantages affecting epidemic severity, and Descloux et al. suggested a correlation of DENV within-host diversity with disease severity [9, 10]. iSNVs have also been used for more precise tracking of viral transmission and spread within outbreaks and epidemics, providing more granular information on transmission chains and possible hotspots of viral dissemination [11, 12].

NGS of within-host viral populations is preferably done directly on clinical samples, such as blood, serum, nasopharyngeal swabs or saliva, cerebro-spinal fluid, and others. However, in many cases, clinical samples may have low viral loads, thus not providing enough genetic material for deep sequencing and complete genome reconstruction with high depth of coverage, requirements that are needed for analyses of low-frequency variants within a sample. This creates bias in some DENV genomic epidemiological investigations as low-viremic individuals are excluded from genomic analysis yet likely play a major role in DENV epidemic dynamics [7, 13].

An alternative approach is to isolate the virus from clinical samples in cell culture and passage it a low number of times (typically less than three passages) in order to amplify the virus thereby allowing an increase in sequencing depth of coverage. Generally, it is thought that isolation and low-passaging does not change the consensus sequence of the virus. For instance, Vasilakis et al. investigated how passaging affects the consensus sequence of DENV, and although they did not compare direct clinical sample sequence to the sequence of the passaged isolates, they showed that passaging of isolates many times (n=5 or n=10) results in change of the virus consensus sequence while low passage number (n=2) does not [14]. They suggested that these changes are most probably not due to a passaging bottleneck, but rather adaptation to cells. Importantly, they also showed that fewer passaging-induced consensus mutations are found in viruses passaged through C6/36 cells *(Aedes albopictus* vector) compared to passaging in human cell lines or vector-human alternating cell lines. Likewise, Chen et al. showed an increasing number of mutations in DENV consensus sequences with the increased number of isolate passages (n=20, 30) [15]. However, no studies to date have compared how the sequence of DENV virus from a direct clinical sample may change due to passaging.

In addition to the lack of knowledge on how DENV isolation from a clinical sample impacts the consensus sequence, there is no information on how the intra-host virus population may change with isolation and passaging. This information is scarce for arboviruses in general. Stapleford *et al.* compared Chikungunya virus (CHIKV) iSNVs in isolate samples, with single passage in mammalian or vector cell lines, with iSNVs in viruses from direct clinical samples [16]. They found that the high frequency iSNVs were maintained over passaging. However, they also noted a decrease in overall diversity, with more diversity maintained when passaging one time through mammalian cells rather than vector cells. No such studies have been performed on DENV.

Because of the importance of accurate DENV consensus genomic data from a range of dengue case viral loads, as well as accurate estimations of the intra-host populations and minor variants, we sought to compare DENV consensus sequences from direct clinical samples (sera) to low-passage isolates derived from the same samples. In addition, we compare iSNV (minor variant) frequencies in DENV sequencing both in direct samples and their corresponding isolate genomes. This provides information on the possible consensus changes low-passaging of DENV from direct clinical samples might induce, as well as any changes in the frequency of intra-host population variants. Since low-passaged DENV isolates are frequently and increasingly used in investigations of DENV diversity and spread, this study provides important insight into the effect of passaging on the DENV genome sequence.

## MATERIALS AND METHODS

### Specimen collection and viral passaging

Eleven DENV1 culture-positive sera and their corresponding isolates, and 14 DENV2 culture-positive sera and their corresponding isolates, were received from the Armed Forces Research Institute of Medical Sciences (AFRIMS). These specimens were collected between 2012 and 2015 from a prospective cohort study of incident dengue illness set in Cebu, the Philippines (WRAIR #1844). The isolates had been passaged as follows: undiluted plasma/serum (0.34 μl/mosquito) was inoculated in 10-20 *Toxorhynchitis splendens* mosquitoes. Amplification of virus in mosquitoes was assessed by an immunofluorescence assay (IFA) assay on head squash preparations. The bodies of IFA-positive mosquitoes were macerated and inoculated onto C6/36 cells. Following one passage, the virus was serotyped, aliquoted, and stored at −70°C or below.

### Virus sequencing

200ul of each clinical specimen and corresponding cultured isolate was used for RNA extraction with QIAamp Viral RNA Mini Kit (Qiagen, MD, USA). Full DENV genomes were amplified via Access Array (AA) system (Fluidigm Corporation, CA, USA) using 48 pairs of in-house DENV serotype-specific primers and the SuperScript III One-Step RT-PCR system with Platinum® Taq High Fidelity polymerase (ThermoFisher Scientific, MA, USA), followed by cDNA purification using Agencourt AMPure XP beads (Beckman Coulter, CA, USA). The PCR products were analyzed using High Sensitivity DNA D5000 tapes (Agilent Technologies, CA, USA) on the Agilent Tapestation 4200 System (Agilent Technologies) to check cDNA quality and quantity. The next-generation sequencing (NGS) libraries were prepared using QIAseq FX DNA Library kit (Qiagen) following the manufacturer’s instructions. 50ng per sample of amplified amplicon was used as input DNA for library preparation. The input DNA was fragmented for 15 minutes and amplified 6 cycles. The purified indexed libraries were quantified with Agilent Tapestation using DNA D5000 tapes (Agilent). Equal molar quantities of libraries were pooled based on the Tapestation data. Pooled libraries were denatured and diluted to a final loading concentration of 11.5pM and loaded onto the Miseq sequencing reagent cartridge from Miseq Reagent kit v3, 600 cycles, (illumina, CA, USA) for sequencing.

### Virus genome assembly and variant calling

DENV reads from each sample were mapped to one of four references: one DENV1 reference (AB204803) and three DENV2 references (KU509277, KM279601 and KU517847), which were determined by first mapping the sequenced samples to a concatenated reference file containing numerous DENV1 and DENV2 genomes with location listed as Philippines and year 2012 to 2015 from GenBank. Using mapping output to support the best-fitting reference for each sample, the sequenced samples were mapped again individually to one of the four references. For the mapping and analysis, we used an in-house pipeline, NGS_Mapper that includes the variant caller LoFreq [17, 18]. Briefly, the default setting of 5/95 was used for the NGS_Mapper calling of intra-host nucleotide variants, allowing for analyses of variant frequencies per nucleotide position down to 5%. The minimum nucleotide Phred score threshold was set to 30. This means that any site containing a variant nucleotide of quality score greater than 30 and present at, or greater than, 5% would be annotated as an ambiguous position. In addition, the minimum depth of coverage to detect a variant at 5% was set to 500X. The consensus genomes were quality checked and manually curated using IGV and Geneious version R10, as well as each sample’s VCF file for statistical support [19–21]. This post-assembly cleaning process of the final consensus was implemented to ensure removal of any nucleotide variants present due to primer-induced error, certain types of sequencing error, and strand bias higher than what was observed in confident calls from each genome, respectively. In addition, minor nucleotide variants that were consistently present at either ends of the reads were manually removed from the analyses, as these have previously been shown to be spurious [22]. All genomes have been submitted to GenBank under accession numbers MT832032-MT832080.

## RESULTS

Full genomes were obtained for all 11 DENV1 viruses derived from the direct clinical samples and for 10 of their corresponding virus isolates *(T.splendens* inoculation followed by one C6/36 passage). Full genomes were obtained from 14 DENV2 viruses derived from the direct clinical samples and from all of their corresponding virus isolates *(T.splendens* inoculation followed by one C6/36 passage). The overall depth of genome coverage for all obtained sequences was >1000x throughout the genome, and positions with intra-host Single Nucleotide Variants (iSNVs) were scanned for. Variants present at a frequency of 5% or higher were called based on criteria described in Materials and Methods.

### DENV1 consensus and iSNVs in direct sample versus corresponding culture

Of 22 DENV1 samples (11 direct-culture pairs), one genome, from the AB204803_D1_11 cultured virus, was not assembled. All except one direct-culture pair had identical consensus genomes (Table 1). One genome, from the AB204803_D1_9 cultured virus, contained an unusual number of iSNV positions and was deemed an outlier, pointing to possible contamination of this sample (Table 1). Thus, nine DENV1 samples were used for DENV1 iSNV comparisons between viruses sequenced directly from clinical samples and their corresponding cultured isolates. Three of the samples did not have any iSNV positions in the genomes derived from either the direct sample or the cultured virus (Table 1). For genomes from direct samples, the average number of iSNV positions was 1 (range 0-3) per genome (Table 1). For genomes from their corresponding isolates, the average number of iSNV positions was 1.1 (range 0-5) per genome (Table 1) and it did not differ significantly from the number of iSNV positions in the direct samples (*p*=0.36, f-test). Out of a total of nine positions with iSNVs in the clinical DENV1 samples, eight were also present in their isolates (89%). Isolate samples had a total of three iSNV positions that were not found in the clinical samples of DENV1.

**Table 1.** Number of consensus differences and positions with iSNVs per DENV1 genome from direct samples and their corresponding isolates.

| Sample name | iSNVs Direct sample | iSNVs Isolate | Consensus differences<sup>a</sup> |
| --- | --- | --- | --- |
| AB204803_D1_1 | 0 | 0 | 0 |
| AB204803_D1_2 | 3 | 5 | 0 |
| AB204803_D1_3 | 1 | 1 | 0 |
| AB204803_D1_4 | 1 | 0 | 0 |
| AB204803_D1_5 | 0 | 0 | 0 |
| AB204803_D1_6 | 0 | 1 | 0 |
| AB204803_D1_7 | 1 | 1 | 0 |
| AB204803_D1_8 | 3 | 3 | 0 |
| AB204803_D1_9 | 0 | 19<sup>b</sup> | 0 |
| AB204803_D1_10 | 0 | 0 | 1 |
| AB204803_D1_11 | 0 | NA | NA |

<sup>a</sup>Not including iSNV positions
<sup>b</sup>Outlier sample
NA – Not assembled

We compared the positions with iSNVs, and variant frequencies within those positions, between the genomes from DENV 1 direct samples and their corresponding cultures (Table 2). We found that many iSNV positions overlapped between the direct sample and culture viral populations. In total, out of 12 DENV1 positions that contained iSNVs, eight contained variants at similar frequencies in the two sample types (Table 2). In four of the positions, a minor variant present at the frequency of ≥5% was found exclusively in either the direct sample or the isolate genome. We cannot say with certainty that these minor variants did not exist in the variant-negative genomes at a frequency lower than 5%, since that was our threshold cutoff for confident variant determination. We also calculated differences in iSNV minor variant frequencies between DENV1 direct samples and cultured samples. The average difference in minor variant frequency was 6.8% (95CI 3.2%) and ranged from 0-18%. Given the length of the DENV1 Coding Sequence (CDS) of 10,179 nucleotides, the mean per position (with variant cutoff at 5%) nucleotide call concordance (including called iSNVs) between direct samples and their cultured viruses was 99.99%.

**Table 2.**
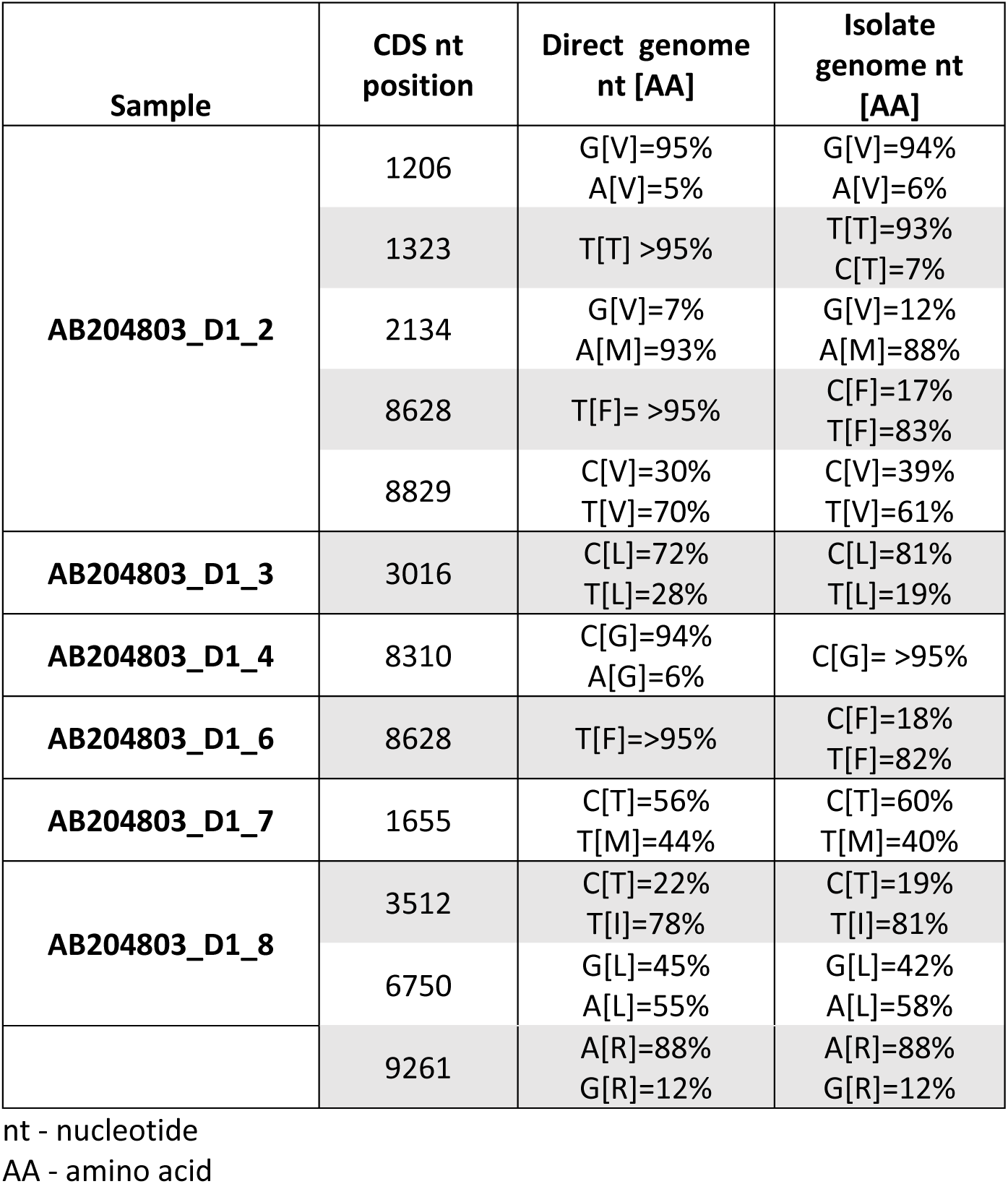
DENV1 iSNV CDS positions and their variant frequencies for samples mapped to reference AB204803.

| Sample | CDS nt position | Direct genome nt [AA] | Isolate genome nt [AA] |
| --- | --- | --- | --- |
| AB204803_D1_2 | 1206 | G[V]=95%<br>A[V]=5% | G[V]=94%<br>A[V]=6% |
|  | 1323 | T[T] >95% | T[T]=93%<br>C[T]=7% |
|  | 2134 | G[V]=7%<br>A[M]=93% | G[V]=12%<br>A[M]=88% |
|  | 8628 | T[F] = >95% | C[F]=17%<br>T[F]=83% |
|  | 8829 | C[V]=30%<br>T[V]=70% | C[V]=39%<br>T[V]=61% |
| AB204803_D1_3 | 3016 | C[L]=72%<br>T[L]=28% | C[L]=81%<br>T[L]=19% |
| AB204803_D1_4 | 8310 | C[G]=94%<br>A[G]=6% | C[G] = >95% |
| AB204803_D1_6 | 8628 | T[F]=>95% | C[F]=18%<br>T[F]=82% |
| AB204803_D1_7 | 1655 | C[T]=56%<br>T[M]=44% | C[T]=60%<br>T[M]=40% |
| AB204803_D1_8 | 3512 | C[T]=22%<br>T[I]=78% | C[T]=19%<br>T[I]=81% |
|  | 6750 | G[L]=45%<br>A[L]=55% | G[L]=42%<br>A[L]=58% |
|  | 9261 | A[R]=88%<br>G[R]=12% | A[R]=88%<br>G[R]=12% |

nt - nucleotide
AA - amino acid

### DENV2 consensus and iSNVs in direct sample versus corresponding culture

Of 28 DENV2 samples (14 direct-culture pairs), all genomes were available for consensus and iSNV comparisons between clinical samples and their corresponding cultured viruses. There were 0-1 consensus genome differences found between the sample pairs (Table 3). Two of the samples did not have any iSNVs in either the direct sample or the corresponding cultured virus genomes (Table 3). For DENV2 genomes from direct samples, the average number of iSNV positions was 2.2 (range 0-5) per genome. For genomes from their corresponding isolates, the average number of iSNV positions was 2.4 (range 0-6) per genome (Table 3). The number of iSNV positions did not differ significantly between direct samples and their isolates (*p*=0.64, f-test). Out of a total of 31 iSNV positions in DENV2 clinical samples, 20 were also present in their isolates (65%). Isolate samples had a total of 14 iSNV positions that were not found in the clinical samples of DENV2.

**Table 3.** Number of consensus differences and positions with iSNVs per DENV2 genome from direct samples and their corresponding isolates.

| Sample name | iSNVs Direct sample | iSNVs Isolate | Consensus differences<sup>a</sup> |
| --- | --- | --- | --- |
| KU509277_D2_1 | 2 | 2 | 0 |
| KU509277_D2_2 | 3 | 3 | 0 |
| KU509277_D2_3 | 2 | 2 | 0 |
| KU517847_D2_1 | 1 | 1 | 0 |
| KM279601_D2_1 | 5 | 6 | 1 |
| KM279601_D2_2 | 1 | 2 | 1 |
| KM279601_D2_3 | 3 | 1 | 1 |
| KM279601_D2_4 | 4 | 3 | 1 |
| KM279601_D2_5 | 3 | 5 | 1 |
| KM279601_D2_6 | 0 | 0 | 0 |
| KM279601_D2_7 | 0 | 0 | 0 |
| KM279601_D2_8 | 2 | 3 | 0 |
| KM279601_D2_9 | 4 | 4 | 0 |
| KM279601_D2_10 | 1 | 2 | 1 |

<sup>a</sup>Not including iSNV positions

We compared the positions with iSNVs, and variant frequencies within those positions, between the genomes from DENV2 direct samples and their corresponding cultures (Table 4 and Table S1). In total, 20 of 45 DENV2 positions contained variants at similar frequencies in the two sample types. As for DENV1, DENV2 positions with a minor variant present at the frequency of ≥5% were found exclusively in either the direct sample or the isolate genome. We cannot with certainty say that these minor variants did not exist in the variant-negative genomes at a frequency lower than 5%, since that was our threshold cutoff. The average DENV2 variant frequency difference between iSNV positions in direct sample versus the corresponding cultured genomes was 9.6% (95CI 2.6%) and ranged from 0-46%. Given the length of the DENV2 CDS of 10,176 nucleotides, the mean per position nucleotide call concordance (with variant cutoff at 5%) between direct samples and their cultured viruses was 99.99%.

**Table 4.** DENV2 iSNV CDS positions and their variant frequencies for samples mapped to references KU509277 or KU517847.

| Sample | CDS nt position | Direct genome | Isolate genome |
| --- | --- | --- | --- |
| KU509277_D2_1 | 4056 | T[A]=69%<br>C[A]=31% | T[A]=72%<br>C[A]=28% |
|  | 8085 | T[A]=69%<br>C[A]=31% | T[A]=76%<br>C[A]=24% |
| KU509277_D2_2 | 1014 | G[K]=69%<br>A[K]=31% | G[K]=86%<br>A[K]=14% |
|  | 2253 | C[R]=11%<br>T[R]=89% | T[R]= >95% |
|  | 2338 | G[V]= >95% | G[V]=93%<br>A[I]=7% |
|  | 4374 | A[S]=90%<br>G[S]=10% | A[S]=94%<br>G[S]=6% |
| KU509277_D2_3 | 1059 | C[R]= >95% | C[R]=94%<br>A[R]=6% |
|  | 1098 | G[Q]=95%<br>A[Q]=5% | G[Q]= >95% |
|  | 8642 | C[T]=82%<br>T[I]=18% | C[T]=82%<br>T[I]=18% |
| KU517847_D2_1 | 4959 | T[I]=7%<br>A[I]=93% | T[I]=12%<br>A[I]=88% |

nt - nucleotide
AA - amino acid

There were no significant differences between DENV1 and DENV2 in either the number of iSNV positions per genome, or in the difference in variant frequencies between the sample pairs (direct and isolate) (*p*=0.36 and *p*=0.13, respectively, f-test).

## DISCUSSION

Genomic epidemiology is increasingly being used in dengue and other viral outbreaks to provide complementary insights into viral transmission and epidemic dynamics [23, 24]. Such information can be used for more precise tracking of viral transmission and locating possible hotspots of viral dissemination. However, consensus genomes provide limited epidemiological inference, particularly when sampled from infected cases closely linked in space and time [7].

To circumvent this problem and increase resolution, intra-host single nucleotide variants (iSNVs) have recently been used for more precise reconstructions of virus transmission chains [9–12]. For such studies, sequencing of viral genomes from clinical samples is ideal, however, the amount of genetic material may not be sufficient to achieve depth that is required for iSNV identification and analysis. Therefore, viruses from clinical samples may be amplified in cell culture, to achieve sufficient nucleic acid yields for primary analysis, as well as to preserve the virus for additional studies. Often, research reviews highlight sequencing of isolates as a limitation to direct sequencing of clinical samples, despite a relative absence of data showing passaged viruses having a different consensus sequence to its clinical parent. In this study we aimed to investigate whether low passaging can change DENV genome consensus sequence, or the minor variant admixture, compared to a direct clinical specimen.

We compared virus from direct clinical samples and their corresponding cultured viruses for both DENV1 and DENV2, and found that the nucleotide call concordance (including called iSNVs with variant cutoff at 5%) was on average 99.99%. In addition, the DENV1 and DENV2 comparison showed that there were no significant differences in the number of iSNV positions between the sample pairs (direct and isolate) (p=0.36 and 0.64, f-test for DENV1 and DENV2, respectively). This suggests there was no increase or decrease in overall diversity following amplification of DENV through *T.splendens* inoculation and one C6/36 passage. Although prior work by Vasilakis et al. has shown that repeated passaging of a viral isolate can change the consensus sequence, their study did not compare how low-passaging changed the consensus sequence compared to a direct clinical sample [14]. Here, we demonstrate that low passaging number does not significantly change the consensus sequence of the virus when compared to a direct clinical specimen. This shows that sequencing from low-passaged isolates a viable alternative to sequencing direct clinical samples. This is particularly important for accommodating dengue cases with low viremia in genomic epidemiological investigations [7].

We found that DENV iSNV frequencies in direct clinical samples were largely preserved after mosquito and C6/36 culture. This has not been examined for DENV before, although one study on CHIKV showed maintenance of high-frequency iSNVs during virus passaging [16]. However, it is important to note that we did observe variant positions with low frequency iSNVs that were exclusive to either direct samples or cultured samples, but we cannot with certainty say that the minor variants are unique to one sample type, because our frequency threshold cutoff was 5%. A deeper sequencing study at the 1% or lower cutoff would provide more granular information to make such a determination, however, variants at such low frequencies are difficult to distinguish from background noise and sequencing/algorithm errors. We showed that the average difference in minor variant frequency at the 5% level between direct clinical samples and their isolate cultures for DENV1 was 6.8% (95Cl ±3.2%) and ranged from 0-18%, and for DENV2 was 9.6% (95Cl ±2.6%) and ranged from 0-46%. Relative preservation of clinical sample iSNV frequencies following single passage of the virus was a surprising finding, and it shows that low virus passage may be used for most iSNV estimations when not enough viral material is available in clinical samples. In this study, the viruses were not passaged more than once, and a low-passage approach is commonly used for virus amplification for genomic surveillance studies [14]. Further studies can be done to investigate the impact of increasing number of passages on DENV consensus sequence and maintenance of high and low frequency variants. In addition, further analyses on other DENV serotypes (DENV3 and DENV4) are needed, as well as confirmation of our results using other *in vitro* passage protocols and samples.

In conclusion, we show that limited culture passaging had minimal effect on the consensus sequence of DENV from direct clinical samples, and that iSNV frequencies were largely maintained during passage. These data provide more confidence in using virus isolates as an alternative approach to deep sequence DENV intra-host viral populations, and mitigate epidemiological sampling limitations previously constrained by dengue case viral load. Our results serve as an important technical reference for DENV genomic epidemiology, virus dispersal tracking and vaccine effectiveness studies.

## ACKNOWLEDGMENTS

This material has been reviewed by the Walter Reed Army Institute of Research. There is no objection to its presentation and/or publication. The opinions or assertions contained herein are the private views of the authors, and are not to be construed as official, or as reflecting true views of the Department of the Army, the Department of Defense or the National Institutes of Health. The investigators have adhered to the policies for protection of human subjects as prescribed in AR 70–25. This work was funded by the Armed Forces Health Surveillance Branch (AFHSB) and its Global Emerging Infections Surveillance (GEIS) Section, FY2019 ProMIS ID P0116_19_WR and the National Institutes of Health grant P01AI034533.

## Notes

### Competing Interest Statement

The authors have declared no competing interest.

